# Ultrastructural Analysis of Extracellular Vesicles and Surface Protrusions from the Parasite *Tritrichomonas foetus*

**DOI:** 10.64898/2026.10.09.758062

**Authors:** Veronica M. Coceres, Abigail Miranda-Magalhães, Regina Celia Bressan Queiroz de Figueiredo, Natalia de Miguel, Antonio Pereira-Neves

## Abstract

*Tritrichomonas foetus* is an extracellular flagellated protist responsible for bovine and feline tritrichomonosis, yet several aspects of its cell biology and pathogenicity remain poorly understood. Here, we evaluated the presence of extracellular vesicles (EVs) and membrane protrusions in *T. foetus* using complementary electron microscopy and immunolabeling approaches. EV-like structures of different sizes were found protruding from the parasite cell body and anterior flagella, and ultrathin sections showed continuity between the membranes of these protrusions and the plasma or flagellar membrane. Multivesicular bodies containing intraluminal vesicles were also observed, and an EV-enriched fraction isolated from conditioned medium contained vesicles with a broad size distribution, consistent with heterogeneous EV populations. Immunofluorescence further showed tetraspanin-1 in intracellular vesicles, flagella, and vesicular structures protruding from the parasite surface. Exposure to bovine preputial epithelial cells resulted in a ninefold increase in the percentage of parasites displaying flagellar cytoneme-like structures, suggesting a possible role for these projections in host-parasite interactions. VPS32, a component of the ESCRT-III machinery, was found in intraluminal vesicles, vesicles protruding from the cell body and flagella, isolated EVs, and flagellar membrane protrusions. In parasites overexpressing VPS32, the number of vesicles per flagellum increased, and the percentage of cells with flagellar cytoneme-like projections was fivefold higher than in control parasites. Pearling-like protrusions containing VPS32 were also observed. Together, these findings provide ultrastructural evidence for heterogeneous EV populations associated with the cell body and flagella of *T. foetus* and suggest that VPS32 could contribute to flagellar EV and membrane protrusion formation.

**IMPORTANCE:** *Tritrichomonas foetus* is a veterinary parasite that infects cattle and cats, but important aspects of its cell surface biology remain poorly understood. In this study, we found membrane-bound vesicular structures associated with both the cell body and flagella, including structures whose membranes were continuous with the parasite membrane. We also observed long flagellar projections that became more frequent after contact with host cells, suggesting that they may participate in parasite-host interactions. In addition, the membrane-remodeling protein VPS32 was found in vesicles and surface projections, and overexpression of this protein was accompanied by greater numbers of flagellar vesicles and cytoneme-like projections. These findings expand the current understanding of *Tritrichomonas foetus* cell biology and identify vesicles and membrane projections as structures of potential relevance to parasite communication and interaction with host cells. Their composition, mechanisms of formation, and biological functions remain to be determined.

## INTRODUCTION

The flagellated protist *Tritrichomonas foetus* (Metamonada, Parabasalia) colonizes the reproductive tract of cattle and the gastrointestinal tract of cats, causing bovine and feline tritrichomonosis, respectively. Bovine tritrichomonosis is a venereal disease associated with reproductive failure and substantial economic losses, with asymptomatic bulls serving as the main reservoir (1,2). In cats, *T. foetus* is an important cause of chronic large-bowel diarrhea and colitis (3). Despite its veterinary importance, several aspects of *T. foetus* cell biology and pathogenicity remain poorly understood (3,4).

Because *T. foetus* is an extracellular parasite, its cell surface, including the flagella, and secreted factors represent important interfaces for parasite-host interactions and survival. Detailed ultrastructural characterization of these structures by electron microscopy can therefore reveal surface features potentially relevant to pathogenesis. In this context, extracellular vesicles (EVs) have emerged as important mediators of intercellular communication in parasitic protists. Studies on the related human pathogen *Trichomonas vaginalis* have shown that EVs released from the parasite surface participate in parasite-parasite and parasite-host interactions, influencing adherence to epithelial cells and host immune responses (5–9). EVs are membrane-enclosed, non-replicative particles released by cells (10). The major EV subtypes include microvesicles (MVs), which bud outward from the plasma membrane, and exosomes, which originate as intraluminal vesicles (ILVs) within multivesicular bodies (MVBs) and are released upon fusion of MVBs with the plasma membrane (11). EVs can transport diverse biomolecules and, in parasitic protists, modulate host-cell responses and transfer molecules associated with virulence, drug resistance, and differentiation (10,12–22).

Flagella can also participate in EV-mediated communication. Flagellar or ciliary EVs have been described in mammals, nematodes, and protists, including *Chlamydomonas*, *Trypanosoma brucei*, and *T. vaginalis* (5,22–27). Tetraspanins and components of the Endosomal Sorting Complex Required for Transport (ESCRT) machinery participate in EV biogenesis and release (5,26,28). In particular, the ESCRT-III component VPS32/SNF7 contributes to membrane remodeling and vesicle budding (27,29,30). A VPS32 ortholog has been localized to intracellular vesicles in *T. foetus* (31), but its association with EVs and flagellar membrane protrusions remains unclear.

EVs have not yet been examined in *T. foetus*. Here, we used complementary electron microscopy and immunolabeling approaches to evaluate EV-like structures associated with the cell body and flagella and to examine the ultrastructural morphology of an EV-enriched fraction. We also analyzed the subcellular localization of tetraspanin-1, the ultrastructural localization of VPS32, and the occurrence of flagellar cytoneme-like structures during host-cell interaction and following VPS32 overexpression. Together, these analyses provide new ultrastructural information on EV-like structures and membrane protrusions in *T. foetus*.

## RESULTS

### *T. foetus* releases extracellular vesicles (EVs)

Wild-type parasites analyzed by SEM showed ellipsoidal or rounded EV-like structures ranging from 60 to 350 nm in diameter, sparsely distributed on the parasite surface or forming small clusters of approximately 10 vesicles, predominantly near the posterior third of the cell body (Fig. 1A) or the flagellar emergence region (Fig. 1B). Quantitative analysis showed that 7.8% ± 2.2% of the *T. foetus* population exhibited this morphology. A distinct phenotype characterized by numerous (>20) EV-like structures, measuring 148–590 nm in diameter and distributed relatively uniformly over the parasite surface, was also observed (Fig. 1C). This phenotype was found in 3.2% ± 1.4% of the parasite population. At higher magnification, MV-like structures 300–480 nm wide were also observed protruding from the plasma membrane (Fig. 1D), suggesting that this EV population might be released by the parasite. Some of these structures were bifurcated (Fig. 1D).

**Figure 1.**
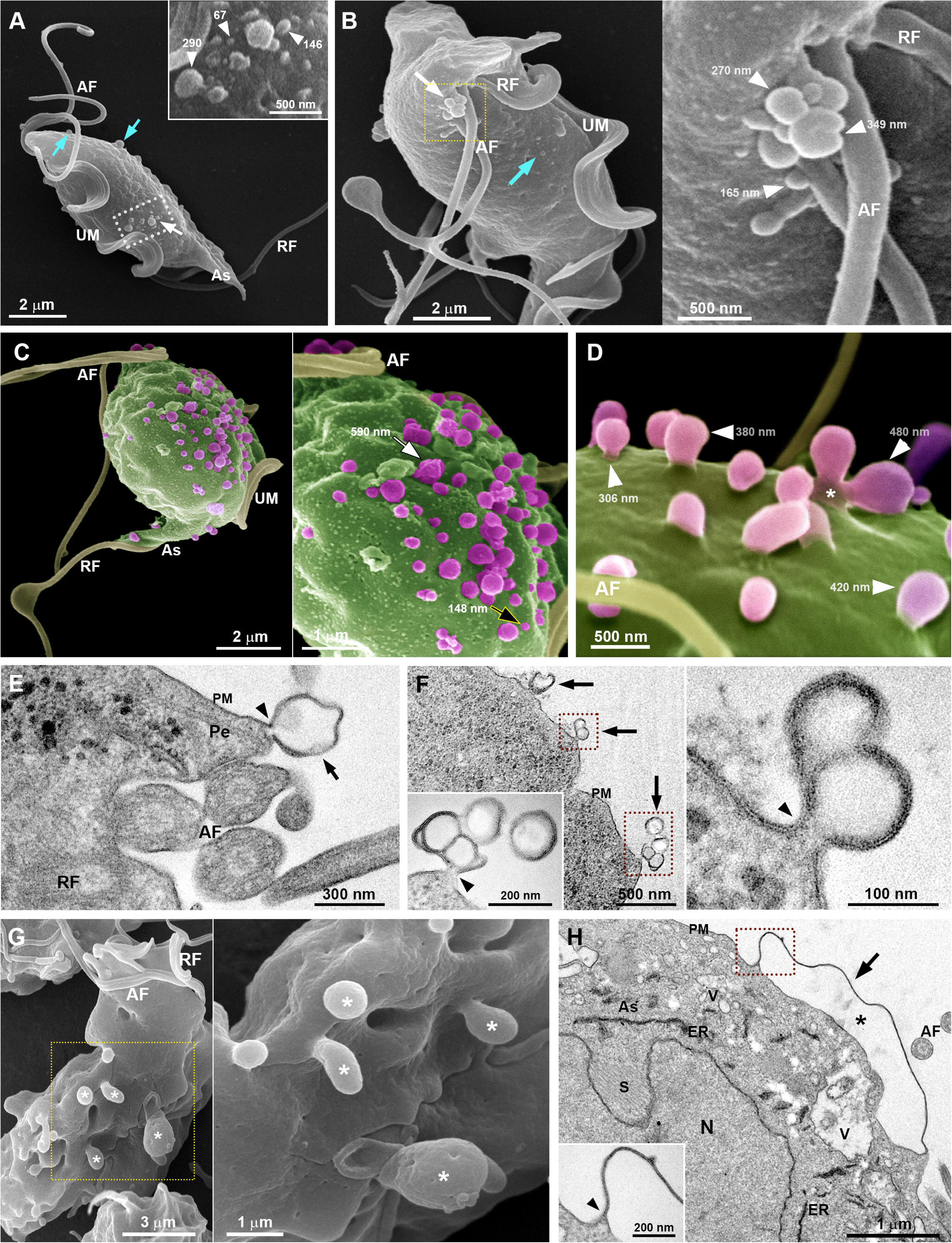
*T. foetus* releases microvesicle-like structures (MVs). A-C. Representative SEM images of MVs on the surface of parasites under growth axenic conditions. The parasites display a pear-shaped cell body, three anterior flagella (AF) and one recurrent flagellum (RF) associated with an undulating membrane (UM). As, axostyle. **A-B**. General and detailed views of parasites showing ellipsoidal or rounded MVs sparsely distributed on the cell surface (blue arrows), together with MVs clusters (white arrows and insets) near the posterior third of the cell body **(A)** or close to the flagellar emergence region **(B).** Representative diameter measurements in nanometers (nm) are indicated (white arrowheads). **C.** General and detailed views of a parasite exhibiting numerous MVs (magenta) on the cell surface (green), ranging from 148 nm (black arrow) to 590 nm (white arrow) in diameter and relatively uniformly distributed on the surface. **D.** Close view of vesicles (magenta), with diameters ranging from 306 nm to 480 nm (white arrowheads), protruding from the cell surface of a parasite (green). Notice that one vesicle has a bifurcated appearance (*). **E-F**. Representative TEM of ultrathin sections of MVs (black arrows) protruding from the anterior region near the pelta (Pe) (**E**) and from the cell body (**F**). The membranes of these protrusions are continuous with the plasma membrane (PM, black arrowheads), and their lumens are electron-lucent. Some vesicles exhibit a bifurcated or bilobed appearance (**F**, insets). **G.** General and detailed SEM images of a parasite showing vesicular structures larger than 1 µm (*) protruding from the surface. **H.** TEM image of an ultrathin section showing a large vesicular protrusion (black arrow) continuous with the plasma membrane (PM, black arrowhead) and with an electron-lucent lumen (*). The presence of a spindle (S) indicates that the parasite is undergoing cell division. N, nucleus; ER, endoplasmic reticulum; V, intracellular vesicles.

TEM confirmed and extended these findings. Ultrathin sections revealed MV-like structures protruding from the anterior region near the pelta, a microtubular structure supporting the flagellar canal (Fig. 1E), and from the cell body (Fig. 1F). These membrane-bound protrusions were continuous with the plasma membrane and displayed an electron-lucent lumen without ribosomes or other typical cytosolic structures. Some vesicles exhibited a bifurcated or bilobed appearance (Fig. 1F).

Vesicular protrusions larger than 1 µm in diameter were also observed on the parasite surface by SEM (Fig. 1G). This phenotype was found in 1.2% ± 0.6% of parasites. In ultrathin sections, these large vesicles were continuous with the plasma membrane and displayed an electron-lucent lumen (Fig. 1H). They were also found in parasites undergoing cell division, as indicated by the presence of the mitotic spindle (Fig. 1H).

TEM also revealed classical ILVs within MVBs in *T. foetus* (Fig. 2A), consistent with the possible presence of exosome-like vesicles. An EV-enriched fraction isolated from conditioned medium contained vesicles of ∼30–100 nm, similar in size and shape to mammalian exosomes, as well as larger vesicles of variable shapes and sizes resembling MVs (Fig. 2B, C). Double-membrane-bound vesicles were also observed. Quantitative analysis of approximately 500 vesicles per sample revealed a broad, right-skewed size distribution, with the highest frequencies between 100 and 199 nm and a progressive decrease above 200 nm; vesicles smaller than 100 nm were also found (Fig. 2D).

**Figure 2.**
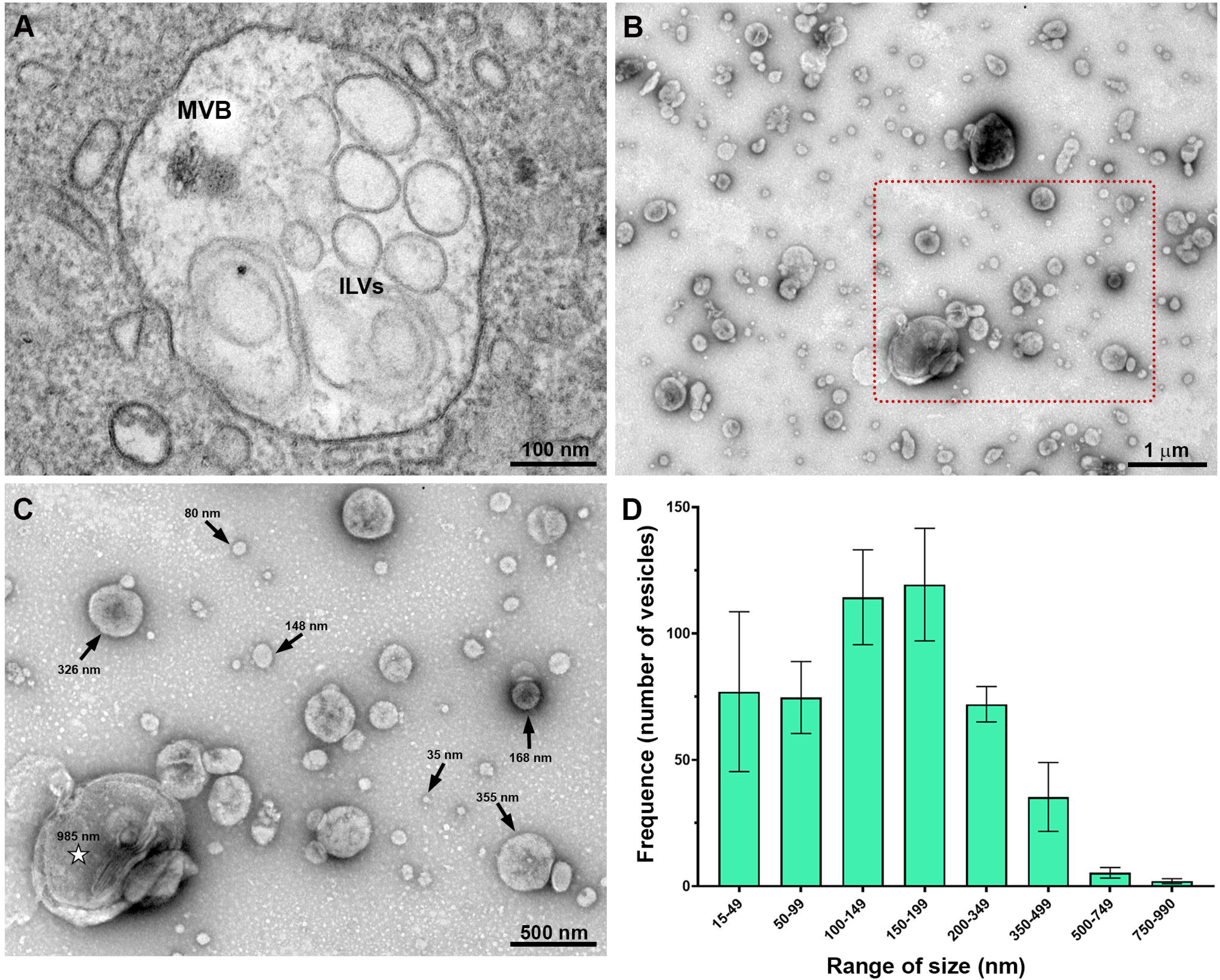
The presence of MVBs, ILVs, and EVs in *T. foetus.* **A.** Ultrathin section of intraluminal vesicles (ILVs) inside a multivesicular body (MVB). **B-C.** Representative TEM images of negatively stained EV-enriched samples showing general (**B**) and detailed (**C**) views of vesicles with different shapes and sizes. (**C**) is representative of the imaging conditions used for the quantitative analysis shown in **(D)**. Diameter measurements (arrows) are indicated in nanometers (nm), and the star marks a double-membrane-bound vesicle. **(D)** Frequency distribution of vesicles according to diameter. Approximately 500 vesicles per sample were measured from randomly acquired TEM images, and diameter values were grouped into predefined intervals. Data represent the mean ± standard deviation of three independent experiments performed in duplicate.

Because TSP1 has been identified in the *T. vaginalis* exosome proteome (6), we investigated its localization in *T. foetus*. Immunolabeling of TfTSP1-HA-transfected parasites showed TSP1 on the cell surface, flagella, and intracellular vesicles, as well as in MV-like structures and large vesicles (>1 µm) protruding from the parasite surface (Fig. 3). Together, these findings support the presence of heterogeneous EV populations associated with *T. foetus* under regular growth conditions.

**Figure 3.**
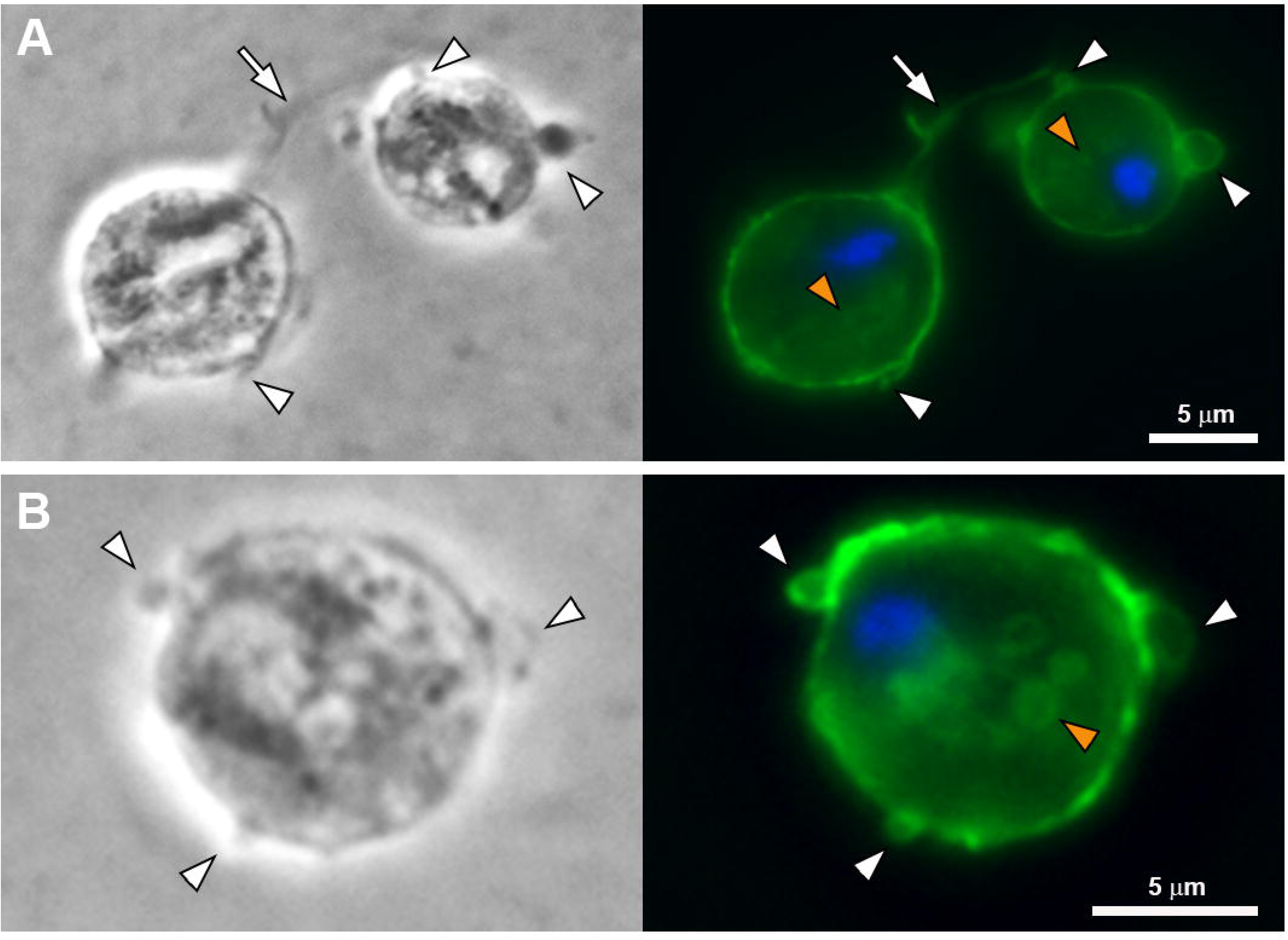
**Subcellular localization of TSP1 protein in *T. foetus*. A-B**. Two representative images of parasites expressing a C-terminal hemagglutinin (HA) tagged version of TfTSP1, incubated with a monoclonal anti-HA antibody, and analyzed using fluorescence microscopy (green). Left-hand panels: Phase contrast microscopy; Right-hand panels: fluorescence. The white arrowheads indicate the localization of TSP1 in MV-like structures and large vesicles (> 1µm) protruding from the parasite’s surface. Positive labelling is also noticed in intracellular vesicles (orange arrowheads) and on the flagella surface (arrow). The nucleus (blue) is stained with 4′,6′-diamidino-2-phenylindole (DAPI).

### MV-like structures can bud from *T. foetus* flagella

We next examined the flagellar surface of *T. foetus* using complementary electron microscopy approaches (Fig. 4). SEM revealed spherical MV-like structures distributed along the anterior flagella, ranging from approximately 57 to 330 nm in diameter (Fig. 4A). This feature was observed in 15.0% ± 3.6% of the cells (n = 500 parasites per sample). We also observed other structures protruding from the flagellar surface, with variable shapes, lengths, and thicknesses/diameters >350 nm (Fig. 4B). These protrusions often displayed a basal constriction (Fig. 4A-B) suggestive of a possible scission site.

**Figure 4.**
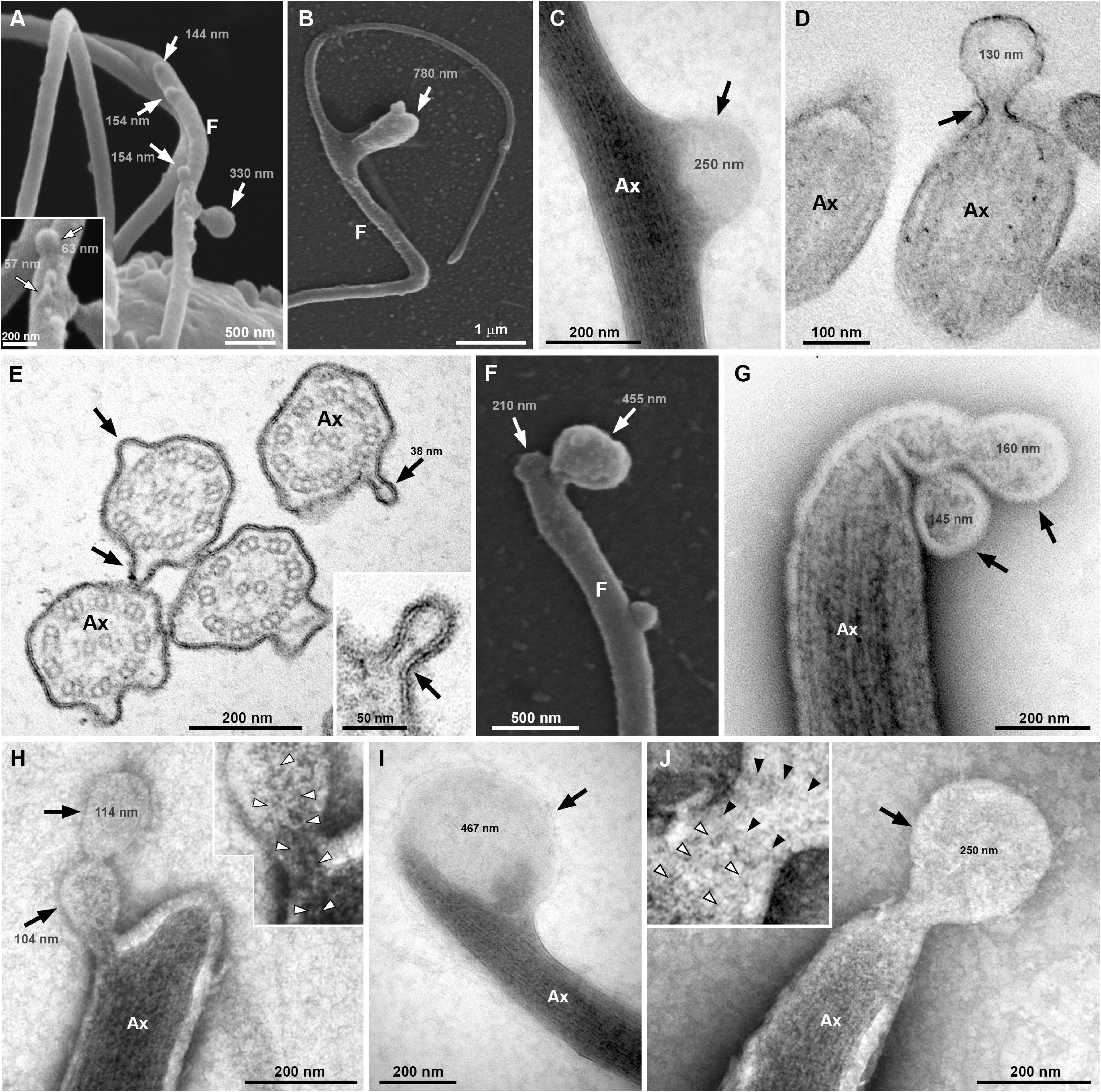
EVs protrude from the anterior flagella of *T. foetus*. A-B. Representative SEM images. **A.** MVs (arrows), ranging from 57 to 330 nm in diameter, protrude along the entire length of the flagellar surface (F). Inset, higher magnification of vesicles smaller than 100 nm protruding from the flagella. **B.** A large protrusion (arrow), approximately 780 nm in diameter, is seen protruding from the flagellar surface (F). **C-E.** Representative TEM micrographs. **C.** Negative staining of an EV approximately 250 nm in diameter (arrow) protruding from the flagellar membrane. **D-E.** Oblique (D) and transverse (E) ultrathin sections showing EVs of varying diameters (arrows) protruding from the flagella. The axoneme (Ax) is tightly surrounded by the flagellar membrane, which is continuous with the EV membrane. The EV lumen is electron-lucent**. F-J.** EVs (arrows), with different diameters (numbers in nm), are seen protruding from the tips of the flagella**. F.** SEM**. G-J.** Negative staining for TEM. Notice that EVs can exhibit a bifurcated (F-G), bilobed (H), or single-vesicle appearance (I-J)**. H, J.** Insets: higher magnification of the constricted region at the base of the vesicle. Notice the presence of fine filamentous structures extending from the flagella toward the vesicles (white arrowheads). A thin septum-like structure is also observed (black arrowheads). Ax, axoneme.

TEM showed continuity between the membrane of MV-like structures and the flagellar membrane (Fig. 4C-E), supporting their origin from the flagellar surface. Their lumen was electron-lucent. Ultrathin sections also revealed MV-like structures smaller than 57 nm, with a mean diameter of 38 nm, protruding from the flagellar membrane (Fig. 4E). Membrane-bound MV-like structures measuring 100–470 nm were also observed at the tips of anterior flagella (Fig. 4F-J), with bifurcated, bilobed, or single-vesicle appearances. Thin filaments 2–5 nm wide extended from the flagella toward some vesicles (Fig. 4H, J), and a thin septum-like structure was observed in the neck connecting the flagellum to a budding vesicle (Fig. 4J), suggestive of a possible scission site. MV-like structures were not observed on the recurrent flagellum or undulating membrane. Together, these observations suggest that the anterior flagella may contribute to EV release.

### Cytoneme-like structures increase during the *T. foetus*-host cell interaction process

To evaluate whether cytoneme-like structures might be associated with host-cell interaction, *T. foetus* parasites were co-incubated with bovine preputial epithelial cells (PECs) and analyzed by SEM (Fig. 5). After host-cell contact, cytoneme-like structures arising from the flagellar base were observed connecting two or more adjacent parasites (Fig. 5A). Cytonemes also emerged from the tips of elongated recurrent flagella (>20 µm) and contacted the PEC surface (Fig. 5B). A similarly elongated recurrent flagellum was observed under axenic conditions (Fig. S1), showing that it can reach comparable lengths in the absence of host cells. Bulbous structures protruding from the recurrent flagellum were also found in contact with the host-cell surface (Fig. 5C).

**Figure 5.**
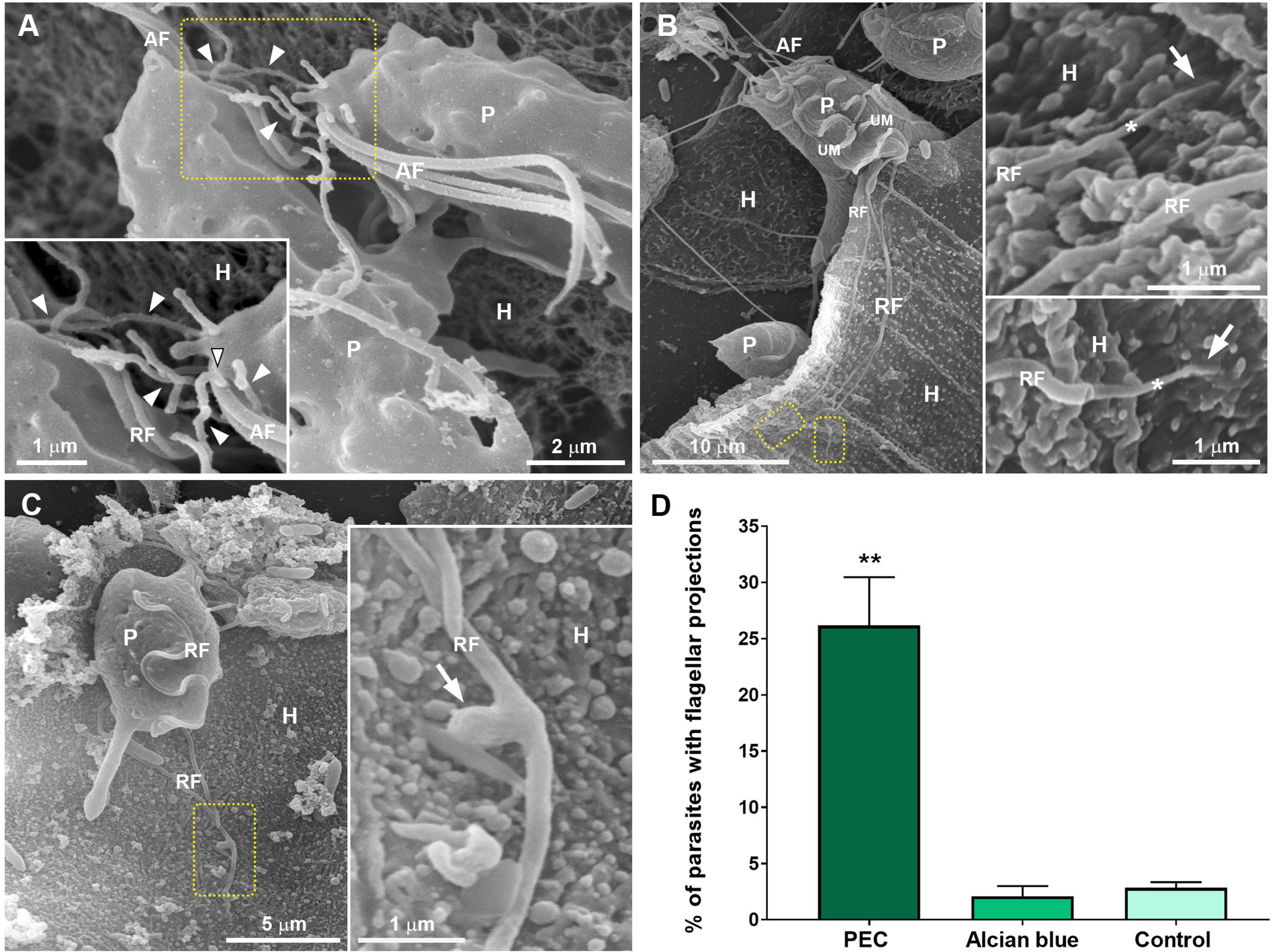
***T. foetus* flagellar cytoneme-like structures increase during host-cell interaction. A-C**. Representative SEM images of *T. foetus* during host-cell interaction. **A.** Cytoneme-like projections (arrowheads) arising from the base of the anterior flagella (AF) are observed connecting two adjacent parasites (P). **B.** Cytoneme-like projections emerging from the tip of an elongated recurrent flagellum (*) are observed in contact with the PEC surface (arrows). In **C**, a bulbous projection (arrow) protruding from the recurrent flagellum (RF) is observed in close contact with the host cell (H). **D.** Quantification of the percentage of *T. foetus* with flagellar cytoneme-like projections during host-cell interaction. Three independent experiments performed in duplicate were analyzed, and 500 parasites were randomly counted per sample using SEM. The data are expressed as the mean percentage of parasites ± SD. In the control experiments, parasites incubated in PBS in the absence of host cells were analyzed. Parasites adhered to Alcian blue precationized coverslips were used as a control for nonspecific adherence. The percentage of parasites with flagellar cytonemes increased after host-cell exposure compared with both controls (PBS and Alcian blue). **p < 0.01 compared with both control groups using the Kruskal–Wallis test followed by Dunn’s multiple comparisons test.

Parasites incubated in PBS and parasites adhered to Alcian blue-coated coverslips were used as nonadherent and nonspecific-adhesion controls, respectively. PEC exposure resulted in a ninefold increase in the percentage of parasites displaying flagellar cytoneme-like structures compared with both controls (Fig. 5D), indicating that this increase was associated with host-cell exposure rather than nonspecific adhesion. These findings suggest a possible role for flagellar cytonemes in cell-to-cell interactions during pathogenesis.

### Exogenous expression of VPS32 increases flagellar EV-like structures in *T. foetus*

VPS32 has been shown to contribute to the biogenesis of EVs in *T. vaginalis* (39). To determine whether VPS32 localizes to compartments associated with EVs in *T. foetus*, parasites expressing HA-tagged full-length TfVPS32 were analyzed by immunoelectron microscopy (Fig. 6). VPS32 was found in ILVs inside MVBs (Fig. 6A), in vesicles located in the extracellular environment next to the plasma membrane of the parasites (Fig. 6B), and MVs protruding from the cell surface (Fig. 6C). Immunogold labeling of isolated EVs further revealed VPS32 in both exosome- and ectosome-like vesicles (Fig. 6D, E).

**Figure 6.**
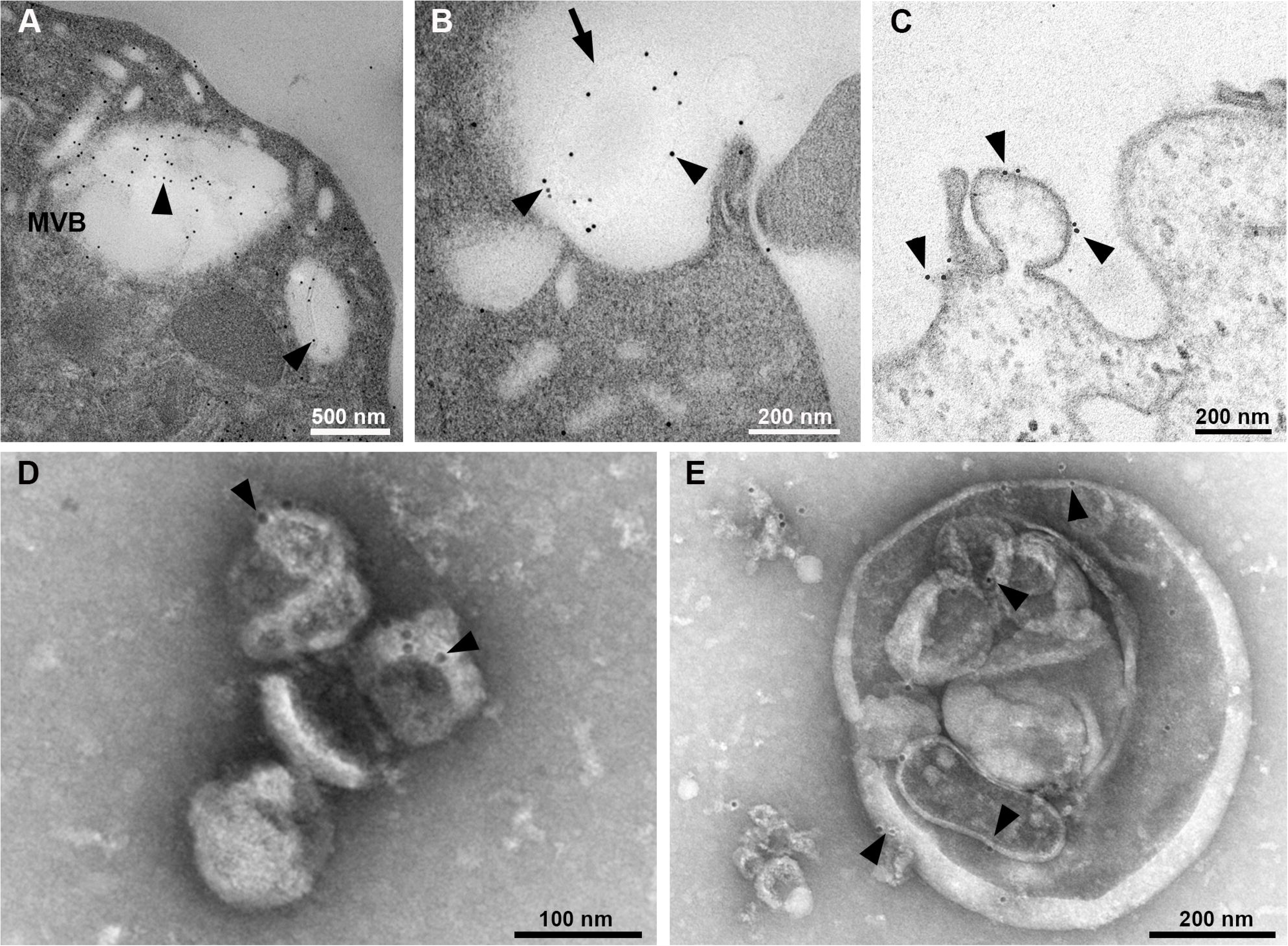
Ultrastructural localization of the TfVPS32 protein. **A**-**C**. Electron micrographs of ultrathin sections of TfVPS32-HA-transfected parasites immunogold-labeled with anti-HA antibodies using post-embedding **(A–B)** and pre-embedding **(C)** labeling techniques. Gold particles (arrowheads) are located on ILVs within a multivesicular body (MVB) (**A**), on the surface of a vesicle located in the extracellular environment (arrow) near the plasma membrane of a parasite (**B**), and on a MV protruding from the parasite surface (**C**). **D**-**E**. Negative staining of isolated EVs from TfVPS32-HA-transfected parasites immunogold-labeled with an anti-HA antibody. Gold particles (arrowheads) are seen on the surface of exosome- (D) and ectosome-like (E) vesicles.

We previously showed that VPS32 localizes to extra-axonemal structures (EASs), and that its overexpression increases these structures in *T. vaginalis* and *T. foetus* (34). Here, TfVPS32 was also found in vesicles protruding along the flagella and from their tips (Fig. 7A). We therefore examined whether VPS32 overexpression correlated with the number of EVs per flagellum. SEM and quantitative analysis revealed more vesicles per flagellum in TfVPS32-transfected than in TfEpNEO parasites (Fig. 7B). In TfEpNEO parasites, 78% of flagella with EVs displayed 1–5 vesicles, whereas 67% of flagella with EVs in TfVPS32-transfected parasites displayed six or more vesicles. These findings suggest that VPS32 could contribute, at least in part, to flagellar EV formation in *T. foetus*.

**Figure 7.**
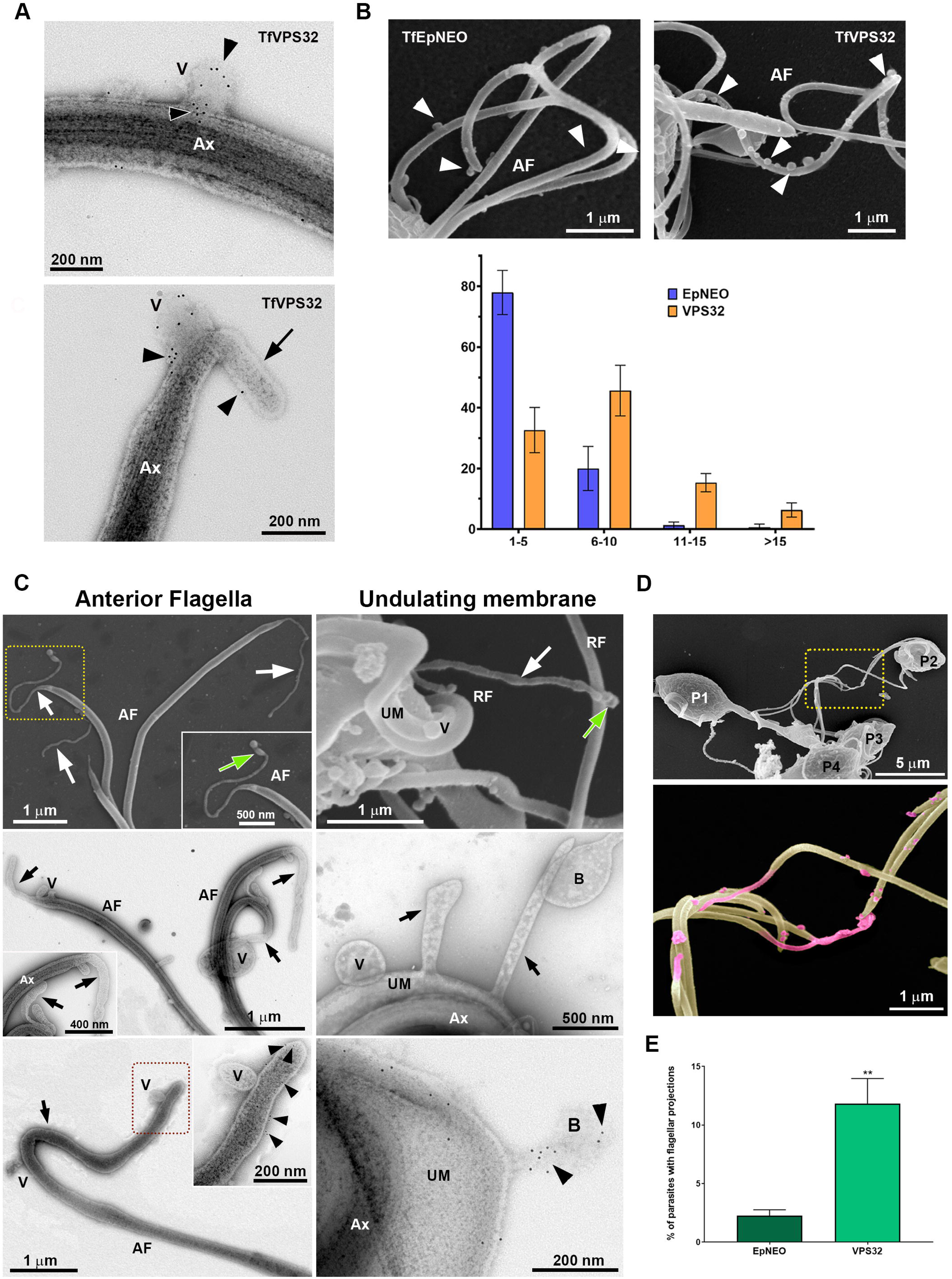
Exogenous expression of VPS32 increases the number of flagellar vesicles and surface protrusions in *T. foetus*. **A**. Negative staining of TfVPS32-HA-transfected parasites immunogold-labeled with anti-HA antibody demonstrates that gold particles (black arrowheads) are localized to vesicles (V) protruding along the flagellum (upper panel) and from its tip (lower panel). Arrow indicates a cytoneme-like projection protruding from the flagellar tip. Ax, axoneme. **B**. Upper panels: representative SEM images of TfEpNEO and TfVPS32 parasites. White arrowheads indicate flagellar EVs. Lower panel: quantitative analysis of the number of EVs per flagellum in TfEpNEO and TfVPS32 parasites. Three independent experiments performed in duplicate were analyzed, and 50 flagella displaying at least one MV-like structure were randomly counted per sample using SEM. The data are expressed as the percentage of flagella within each range of MVs per flagellum ± standard deviation. Notice the higher number of vesicles per flagellum in TfVPS32 parasites compared with TfEpNEO parasites. **C.** Upper panels: representative SEM images. Cytoneme-like structures (white arrows) are seen protruding from the tips of anterior flagella (AF) and the undulating membrane (UM) of VPS32-transfected parasites. Notice vesicle-like structures at the tips of cytoneme-like extensions (green arrows). RF, recurrent flagellum; As, axostyle. Middle panels: negative staining micrographs of TfVPS32 transfected parasites showing cytoneme-like projections (black arrows) protruding from the tips of anterior flagella (AF) and the undulating membrane (UM). Vesicles (V) are seen closely associated with the cytonemes and UM. In the right image, a bulbous protrusion (B) is observed in the extracellular milieu. Ax, axoneme. Lower panels: negative staining of TfVPS32-HA-transfected parasites immunogold-labeled with anti-HA antibody. Gold particles (arrowheads) are seen on the surface of a cytoneme-like projection (black arrow) and a bulbous protrusion (B) protruding from the tip of an anterior flagellum (AF) and the undulating membrane (UM), respectively. Ax, axoneme. **D**. General and detailed SEM views of VPS32-transfected parasites (P) connected to one another by flagellar cytoneme-like projections (magenta). Vesicles closely associated with the cytonemes and flagellar surface (green) are also seen in magenta. **E.** Quantitative analysis of the percentage of *T. foetus* with flagellar cytoneme-like projections in EpNEO and VPS32-transfected parasites. Three independent experiments performed in duplicate were analyzed, and 500 parasites were randomly counted per sample using SEM. The data are expressed as mean ± SD. Notice that the percentage of parasites with flagellar cytoneme-like projections is fivefold higher in TfVPS32 than in TfEpNEO parasites. **p < 0.01, Mann–Whitney U test.

### VPS32 overexpression is associated with increased flagellar surface protrusions in *T. foetus*

Filopodia and other plasma membrane protrusions, such as cytonemes, are known to serve as platforms for EV release, cell communication, and signaling (40). Specifically, these ectosomes are formed by outward shedding, blebbing, pearling, or scission of the plasma membrane or its various protrusions (41). Because ESCRT-III proteins participate in membrane scission, we examined the association between VPS32 and flagellar cytoneme-like protrusions in TfVPS32-overexpressing parasites. SEM and negative-staining TEM revealed various extensions protruding from the tips of anterior flagella and along the undulating membrane (Fig. 7C; upper and middle panels), including bulbous, bead-like, and cytoneme-like projections. Spherical ectosome-like structures were observed at the tips of some cytoneme-like projections (Fig. 7C, upper panels), and vesicles were closely associated with the undulating membrane and these projections. In some cells, tubular or bulbous protrusions displayed constricted connections to the flagellar tips or undulating membrane, suggestive of possible scission sites (Fig. 7C, middle panels). Immunogold labeling showed VPS32 localization in these projections (Fig. 7C, lower panels). TfVPS32-transfected parasites were also observed connecting to one another through flagellar cytoneme-like projections (Fig. 7D). Quantitative analysis showed that the percentage of TfVPS32 parasites displaying flagellar cytoneme-like projections was fivefold higher than that of TfEpNEO parasites (Fig. 7E).

“Pearl necklace-like” protrusions positive for VPS32 were also observed at the tips of flagella in some TfVPS32 parasites (Fig. 8). These structures resembled pearling, a mechanism of MV release involving membrane protrusions (42).

**Figure 8.**
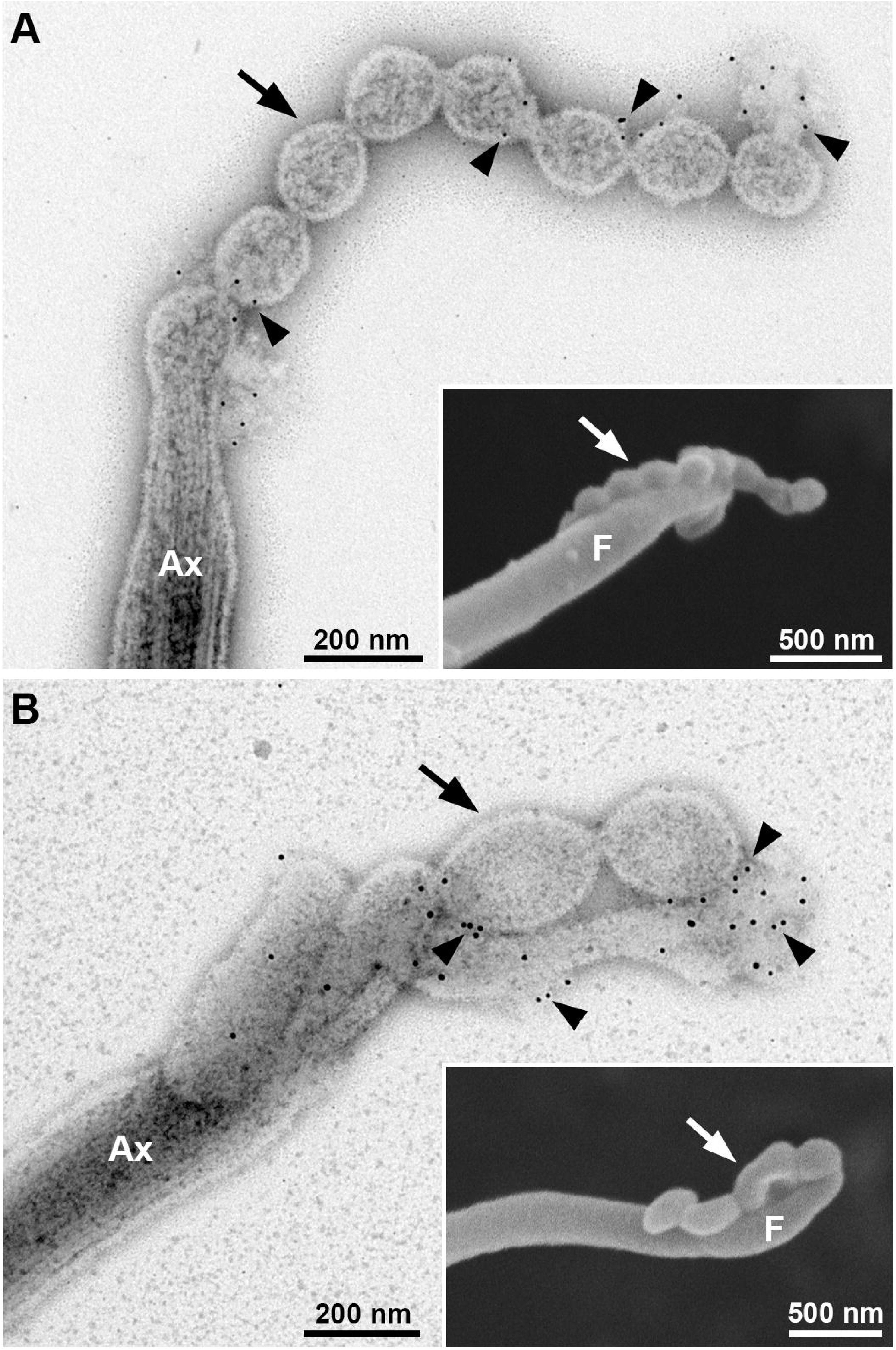
Flagellar pearling protrusions in VPS32-transfected parasites. Representative images of negatively stained TfVPS32-HA-transfected parasites immunogold-labeled with anti-HA antibody and SEM (insets). “Pearl necklace-like” protrusions (arrows) are seen protruding from the tips of flagella (F). Ax, axoneme. Gold particles (arrowheads) are seen on the surface of these projections.

Together, these findings suggest that VPS32 could contribute, at least in part, to the formation of flagellar projections in *T. foetus*.

## DISCUSSION

In recent years, EVs released by protozoan parasites have been implicated in intercellular communication and host manipulation (43). Moreover, EVs can transfer virulence factors, drug-resistance genes, and differentiation factors between parasites (22,44). Among trichomonads, *T. vaginalis* has been reported to release both exosomes and MVs, with the production of these vesicles increasing upon exposure to host cells. This suggests that EVs may play a role in host-parasite interactions (5,6,8,9). In the present study, we identified vesicular structures protruding from the plasma membrane of *T. foetus* and demonstrated the presence of MVBs containing ILVs, consistent with exosome biogenesis. Consistent with these observations, we also isolated an EV-enriched fraction containing vesicles resembling mammalian exosomes in size and morphology, as well as larger vesicles resembling microvesicles. Quantitative analysis further revealed a broad size distribution, with the highest frequencies between 100 and 199 nm followed by a subset of vesicles smaller than 100 nm. This heterogeneous size profile supports the presence of distinct EV populations in *T. foetus*, although vesicle size alone cannot determine their origin. Our results therefore indicate the presence of EV populations consistent with both microvesicles and exosomes in *T. foetus*.

Here, we found that some *T. foetus* vesicle-like structures measured more than 1 µm in diameter, larger than MVs previously reported in other cell models. Vesicles of comparable size have also been observed in *T. vaginalis* (5). Large EVs measuring more than 1 µm have also been described in mammalian cells, including apoptotic bodies, blebbisomes, oncosomes, and exophers (45–49). These structures typically contain cytoplasmic components and, in some cases, organelles (45–49). In contrast, the large vesicular structures observed in *T. foetus* displayed an electron-lucent lumen with no visible cargo in the ultrathin sections examined. To our knowledge, large EV-like structures (> 1 µm) with these ultrastructural features have not yet been described, raising the possibility that they represent a distinct type of vesicular structure in *T. foetus*. Further studies are needed to determine their composition, biogenesis, and biological significance.

Although exosomes and MVs differ in their biogenesis pathways, both processes share molecular components involved in EV generation, such as Endosomal Sorting Complex Required for Transport (ESCRT) proteins and tetraspanins (28,50). Tetraspanins are a superfamily of transmembrane proteins found on the plasma membrane or in endosomal or lysosomal compartments of almost all cell types (51). They are routinely used as markers for small secreted extracellular vesicles (36). Despite extensive research on the protein composition of EVs from parasites, it is still unknown whether the same markers can be used to identify EV populations across different parasite species. Previously, based on the presence of TvTSP1 and TvTSP8 in the exosome and MV proteomes, respectively (5,6), these proteins have been proposed as EV markers in *T. vaginalis*. Accordingly, we found TSP1 in vesicles protruding from the surface of *T. foetus* in parasites transfected with TfTSP1. As previously reported in *T. vaginalis* (52), TfTSP1 is localized to both the cell surface and intracellular vesicles in *T. foetus*. Thus, TSP1 may also serve as an EV marker in this parasite.

Flagella have been shown to mediate intercellular communication through the release of ectosomes (26) which possess a distinct protein composition enriched in proteases, small GTPases, ESCRT and ubiquitinated proteins (27). The ability of cilia to release ectosomes into the extracellular space has been described in a variety of organisms, including *Chlamydomonas* (37,53) *C. elegans* (54), and mammals (23). Trypanosomes produce flagellar EVs that can transfer virulence factors from one parasite to the other, contributing to pathogenesis (22). Moreover, *T. vaginalis* releases flagellar ectosomes that may play an important role in cell communication (5). Interestingly, we also observed that MVs protrude from the flagellar membrane in *T. foetus*. Whether MVs released from the flagella of *T. foetus* differ in content from MVs released from the cell body and whether they may play a role in communication remains to be elucidated.

Components of the ESCRT machinery have been implicated in flagellar ectosome release in other organisms. Specifically, ESCRT-III proteins may play a central role in promoting ectosome budding from the flagellar membrane (27). In addition, VPS32, a member of the ESCRT-III complex, contributes to the biogenesis and cargo sorting of extracellular vesicles in *T. vaginalis* (39). In this context, VPS32 overexpression increased parasite adherence to host cells, supporting a role for this protein in host-*T. vaginalis* interactions (39). Previous studies have shown that VPS32 localizes to the flagellar extra-axonemal structures, and its overexpression increases the formation of these structures in both *Trichomonas vaginalis* and *Tritrichomonas foetus* (34). Moreover, Iriarte et al. (31) reported the localization of VPS32 to intracellular vesicles in *T. foetus*. That study focused on the role of TfVPS32 in cell division and did not investigate its association with EVs or flagellar vesicles. Our findings therefore extend these previous observations by revealing additional TfVPS32-associated compartments in the context of EV formation and release. Here, we demonstrated that TfVPS32 localizes to ILVs within MVBs, extracellular vesicles, and MVs protruding from the cell surface. We also found VPS32 in the isolated EV fraction containing exosome- and ectosome-like vesicles, supporting the hypothesis that VPS32 might be involved in the regulation of EV formation. Additionally, we showed that this protein was also found in vesicles protruding from the flagella, and parasites overexpressing TfVPS32 exhibited an increased number of vesicles per flagellum.

Plasma membranes are fluid and flexible structures, and most cell types possess variable extensions or protrusions supported by the cytoskeleton. The most extensively studied are filopodia, tunnelling nanotubes, microvilli, retraction fibres, lamellipodia, invadopodia, ruffles, and blebs (41). A common feature of plasma membrane protrusions is their ability to act as specialized platforms for EV shedding. Here, we observed cytoneme-like and bead-like projections extending from the flagella of *T. foetus* and confirmed the localization of TfVPS32 to these structures. Parasites overexpressing VPS32 showed a higher percentage of cells displaying these flagellar projections. Host-cell exposure also markedly increased the percentage of parasites with cytoneme-like projections. These structures were observed connecting adjacent parasites as well as extending from the tips of elongated recurrent flagella toward and in contact with the PEC surface, supporting a possible role in cell-to-cell interactions during pathogenesis. In this context, studies in HeLa cells have shown that plasma membrane repair involves recruitment of actin and membrane components to sites of damage, leading to the formation of F-actin-rich filopodia-like protrusions that act as scaffolds for vesicle shedding. ESCRT-III plays a crucial role in membrane scission during the final step in shedding (55), highlighting a relationship between protrusion formation, vesicle release, and the ESCRT machinery. It has been proposed that plasma membrane blebbing and shedding from the tips of filopodia are energetically more favorable than from the flat plasma membrane (41,56), and EVs released from long protrusions extending far from the donor cell body may facilitate EV dispersal within the cellular microenvironment.

Another actin-dependent mechanism for microvesicle release involves membrane nanotubes, which dissociate into EVs through a process known as pearling. This process depends on complex interactions between contractile forces, the underlying cytoskeleton and membrane composition (42). Pearling has been studied in intestinal microvilli, where EVs are shed from the microvillar tips, resulting in the maintenance of microvillar length (57). Notably, we observed pearling-like or vesiculating protrusions at the tips of flagella in *T. foetus*. In this context, microvesicles or ectosomes are formed by outward blebbing, shedding, pearling, or scission of the plasma membrane or its various protrusions, which are platforms for EV release. Pearling is a microtubule-independent mechanism, and it has been reported that localized actin polymerization at the ciliary tip can drive the extrusion of membranous tubes that fragment into EVs (23). A recent study has shown that the glycocalyx can induce plasma membrane instabilities to generate pearled membrane structures and drive the secretion of EVs (58). In our study, TfVPS32 was found in flagellar membrane protrusions, including pearling-like protrusions at the flagellar tips. Thus, we propose that TfVPS32 may participate in the formation and/or scission of these membrane protrusions, potentially contributing to EV release.

The present study explores new aspects of the biology of the parasite *T. foetus*, expanding our understanding of EV formation and release in this parasite.

Taken together, our results provide ultrastructural evidence for heterogeneous EV populations associated with both the cell body and flagella and suggest that TfVPS32 could contribute to flagellar EV and membrane protrusion formation. The increase in cytoneme-like projections during host-cell interaction further points to a possible role for these structures in pathogenesis. Further studies are needed to elucidate the molecular composition and functional roles of these EVs and membrane projections during parasite infection.

## MATERIALS AND METHODS

### Parasite culture and viability assay

*Tritrichomonas foetus* K strain and transfectants were cultured as previously described (34). TfTSP1-HA, TfVPS32-HA, and TfEpNeo transfectants were maintained with 100 μg/ml G418 (Invitrogen). Before each assay, parasite viability was assessed by motility and trypan blue exclusion (0.4% in sterile PBS) using a Neubauer chamber. Only cultures with >95% viability were used.

### Plasmid construction and exogenous protein expression

The full-length TfTSP1 gene was PCR-amplified using the following primer pairs: forward (5’-3’ CATATGATGATTGGAACCTTAAA) and reverse (5’-3’ GGTACCGAATTTTGCAATTCCTT); with *NdeI* and *KpnI* restriction sites engineered, respectively. PCR products were generated using standard procedures and cloned into the Master-Neo-(HA)_2_ plasmid for transfection into *T. foetus*. Electroporation of *T. foetus* K strain was carried out as described previously (32), with 50 μg of circular plasmid DNA. Transfectants were selected with 100 µg/ml G418 (Sigma). The TfVPS32 construct was generated and transfected into *T. foetus* K as previously described (31).

### Electron microscopy

#### Scanning electron microscopy (SEM)

Sample preparation and imaging were performed as previously described (34), except that cells were washed three times with PBS prewarmed to 37 °C and coated with a 20-nm gold-palladium layer before observation.

#### Transmission electron microscopy (TEM). Ultrathin sections

Samples were processed and imaged as previously described (34) *Negative staining:* Parasites were processed as previously described (34), with the following modifications: samples were deposited onto glow-discharged carbon film nickel grids for 5 min at 37 °C, and no permeabilization step was performed before negative staining with 1% aurothioglucose for 5 s. EV samples were fixed in 2.5% glutaraldehyde in 0.1 M cacodylate buffer (pH 7.2), deposited onto glow-discharged carbon film nickel grids and negatively stained as described above.

### Immunogold labeling

Three complementary procedures were used. Negative controls were prepared by omitting the primary antibody and incubating the samples with the gold-labeled secondary antibody only; no labeling was observed under these conditions.

*Postembedding and preembedding labeling.* Samples were processed and immunolabeled as previously described (39), using an anti-HA tag antibody (Invitrogen, 5B1D10) followed by a 10-nm gold-labeled goat anti-mouse IgG antibody (BB International, UK).

*Negative stain labeling.* Parasites and EV samples were settled onto nickel grids as described above and fixed for 1 h at room temperature in 4% paraformaldehyde and 0.5% glutaraldehyde in PEME (100 mM PIPES, 1 mM MgSO₄, 2 mM EGTA, and 0.1 mM EDTA, pH 6.9). After washing, samples were quenched and immunolabeled with anti-HA and 10-nm gold-labeled anti-mouse antibodies as described above, washed with water, negatively stained with 1% aurothioglucose, and observed by TEM.

### Immunofluorescence

TfTSP1-HA-transfected parasites were processed for immunofluorescence as previously described (31). Samples were incubated with anti-HA primary antibody (Covance; 1:500) for 2 h at room temperature, followed by Alexa Fluor-conjugated secondary antibody (Molecular Probes; 1:5,000) for 1 h at room temperature. Adobe Photoshop v. 22.4.3 (Adobe Systems) was used for image processing.

### Isolation of EVs

EV isolation was performed using wild-type, TfEpNEO-, or TfVPS32-transfected *T. foetus* (10^6^ cells/mL). A total of 1 × 10^9^ parasites were washed and incubated for 4 h at 37 °C in 500 mL of serum-free TYM. Conditioned medium was centrifuged (400 × g for 10 min) to remove parasites and cell debris, filtered through a 0.8-μm filter, and centrifuged at 100,000 × g for 90 min to obtain an EV-enriched fraction. The pellet was resuspended in 200 μL cold PBS containing 1× HALT protease inhibitors (Thermo Scientific).

### Parasite-host cell interaction assays

Bovine preputial epithelial cells (PECs), kindly provided by Dr. Maria Aparecida da Gloria Faustino and Dr. Milena Paiva Cavalcanti, were co-incubated with *T. foetus* at a 5:1 parasite:host cell ratio for 30 min at 37 °C, as previously described (34). Parasites incubated in PBS were used as controls, and parasites adhered to 1% Alcian blue-coated coverslips were included as a control for nonspecific adhesion. Samples were analyzed by SEM.

### Quantitative analysis

The percentage of parasites that contain cell body MVs, flagellar MVs- and cytoneme-like structures was determined from counts of 500 parasites randomly selected per sample, using SEM. The number of flagellar MV-like structures per flagellum was determined from counts of 50 flagella displaying at least one MV-like structure, using SEM. For analysis of vesicle size distribution, 10 TEM images per sample, each containing approximately 50 vesicles were randomly acquired at 60,000× magnification. Vesicle diameters were measured using Tecnai Imaging and Analysis (TIA) software (FEI Company). The results are presented as the mean ± standard deviation of three independent experiments performed at least in duplicate. Statistical comparisons were performed, as appropriate, using Student’s *t*-test, one-way ANOVA, Mann-Whitney U test, or Kruskal-Wallis test followed by Dunn’s multiple comparisons test. Statistical analyses were performed using GraphPad Prism v. 7.04 (California, USA). *p* < 0.05 was considered statistically significant.

## ACKNOWLEDGMENTS

We thank Dr. Marlene Benchimol from Universidade do Grande Rio for kindly providing *T. foetus* K strain. We thank Dr. Milena Paiva Cavalcanti and Dr. Maria Aparecida da Gloria Faustino from Instituto Aggeu Magalhães and Faculty of Veterinary Medicine/Rural Federal University of Pernambuco, respectively, for kindly providing PECs. We thank Dr. Karina Saraiva and Dr. Cássia Docena from the Technological Platform Core of the Aggeu Magalhães Institute for their technical support. Ethics approval and consent to participate were not applicable to this study. All data generated or analyzed during the study are included in this article. AMM is a PhD fellow supported by by the Coordenação de Aperfeiçoamento de Pessoal de Nível Superior (CAPES), Brazil. This research received no specific grant from any funding agency in the public, commercial, or not-for-profit sectors. The authors declare that they have no competing interests. VC, AMM, NdM, and APN contributed to the conceptualization of the study. VC, AMM, and APN contributed to the methodology and formal analysis, while VC and AMM conducted the investigation. VC and AMM prepared the original draft of the manuscript, and RCBQF, NdM, and APN contributed to its review and editing. VC, AMM, and APN contributed to visualization. RCBQF, NdM, and APN supervised the study.

## SUPPLEMENTARY FIGURE

Figure S1. Representative SEM image of a parasite maintained under axenic conditions showing an elongated recurrent flagellum (RF) extending approximately 27 µm beyond the cell body. AF, anterior flagella; UM, undulating membrane.

