## Supplementary figures and images for "Ultrastructural Analysis of Extracellular Vesicles and Surface Protrusions from the Parasite *Tritrichomonas foetus*"

### Fig S1

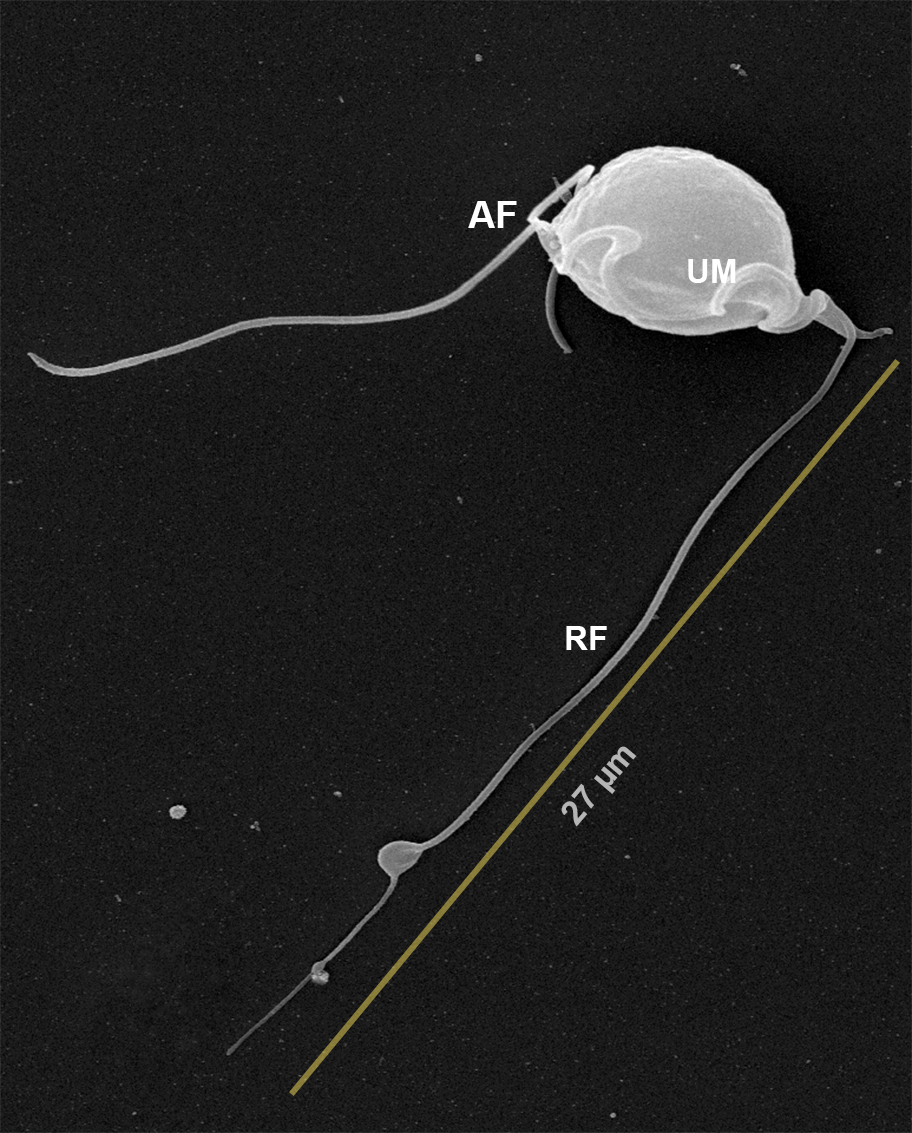
